# An Adaptable Prosthetic Wrist Reduces Subjective Workload

**DOI:** 10.1101/808634

**Authors:** Nathaniel R. Olsen, Jacob A. George, Mark R. Brinton, Michael D. Paskett, David T. Kluger, Troy N. Tully, Christopher C. Duncan, Gregory A. Clark

**Affiliations:** Department of Biomedical Engineering, University of Utah, Salt Lake City, UT; Blackrock Microsystems, Salt Lake City, UT; Department of Computer Science, Westminster College, Salt Lake City, UT; Department of Physical Medicine and Rehabilitation, University of Utah, Salt Lake City, UT

## Abstract

Many presently available prostheses lack a functional wrist. To fill this niche and to better understand the impact a wrist has in prosthetic functionality, we designed a low-cost, adaptable, 3D-printable prosthetic wrist that can be adapted to various prosthetic hands and sockets. The wrist utilizes inexpensive but powerful servo motors to provide simultaneous and proportional control of two degrees of freedom: pronation/supination and flexion/extension or radial/ulnar deviation. Participants used both our wrist and a commercially available wrist (DEKA “LUKE” Arm) to complete a modified version of the clothespin relocation task with and without the wrists enabled. Through use of the NASA Task Load Index we found that both wrists significantly reduced the subjective workload associated with clothespin relocation task (*p* < 0.05). However, we found no significant difference in task completion speed, presumably due to compensation strategies. This inexpensive and adaptable prosthetic wrist can be used by amputees to reduce task workload, or by researchers to further explore the importance of wrist function.

## INTRODUCTION

In 2005, there were an estimated 41,000 upper-limb amputees (excluding partial-hand) in the United States [1]. Amputation results in a life-long struggle with chronic pain, depression and functional disability [1–3]. The current standard of care, often a body-powered hook or myoelectric prosthetic hand, is unsatisfactory, causing up to 50% of amputees to abandon their prostheses [4]. Transradial amputees have expressed their top priorities for useful prostheses (in order of importance): wrist rotation, simultaneous movements, wrist deviation, wrist flexion/extension, increased automaticity [5]. Additional priorities include reduced weight, improved durability, and increased strength [5].

Functional wrist motion is important in reducing biomechanical strain. Without a wrist, amputees are forced to compensate with unnatural movements to complete standard activities of daily living (ADLs) [6–8]. The continual use of these motions causes further damage to the musculoskeletal system over long periods [9,10].

3D-printing has provided new and innovative solutions to address several of the above issues with prosthetics. For example, 3D-printing has considerably lowered the cost and weight of prostheses [11–13], and several 3D-printed prosthetic hands are currently available [14–16]. Traditionally, wrists have been excluded from the design of 3D-printed and commercial prosthetic arms to conserve space and to focus on the implementation of a more dexterous hand [17]. Commercially available prosthetic hands without an active wrist include the Michelangelo [18], i-limb Ultra [19], TASKA [20], and bebionic [21].

Although these commercially available prosthetic hands can be paired with a wrist, few commercial active wrists exist, and none provide more than 1 active degree of freedom [22,23]. There are many dedicated, passive wrists with varying degrees of freedom – such as the bebionic [21] – but these are unnatural since the wrist must be manually adjusted by the intact contralateral hand of the user – at which point, most unilateral tasks could have been completed more efficiently without involving the prosthetic.

To date, the only prosthetic hands fully integrated with an active wrist are the DEKA “LUKE” Arm [24] and the Modular Prosthetic Limb [25]. However, the DEKA “LUKE” Arm is expensive and the Modular Prosthetic Limb is not available to the public. To make it possible for more researchers to study prosthetic wrists, and for amputees to have access to an inexpensive alternative, there is a critical need for an adaptable, inexpensive prosthetic wrist.

Here we describe the development of an active, 3D-printed, inexpensive, and adaptable prosthetic wrist. We show that the implementation of this wrist as well as a commercial wrist does not improve the time it takes to complete tasks but significantly reduces subjective workload in both cases by a factor of about 1.6. This work demonstrates the first standalone, lightweight and 3D-printed prosthetic wrist capable of interfacing with various prosthetic hands and sockets.

## METHODS

### Design Criteria

#### Degrees of Freedom and Length

The human wrist moves in two degrees of freedom: flexion/extension and radial/ulnar deviation. Pronation/supination is frequently attributed to the wrist; however, it is actually a function of the forearm. In prosthetic wrist design, pronation/supination is the most commonly included degree of freedom. Radial/ulnar deviation is included less frequently in wrist designs and is often viewed as the least important of the three degrees of freedom [8].

A transradial amputation occurs across the radial bone, below the elbow but above the wrist. Assuming the average transradial amputation occurs halfway down the forearm, it would be ideal for a wrist to not exceed half the length of the average human forearm. This length, averaged between males and females, is roughly 12.5 cm [26]. By limiting the length of the wrist in this way, the wrist would be more easily integrated with a prosthetic hand without dramatic increases to the length of the prosthetic device.

#### Adaptable

Existing wrists are not easily adaptable to multiple hands. The Ottobock Quick Disconnect Wrist (Otto Bock HealthCare LP, Austin, TX) fits a variety of hands, but prostheses not equipped with the Ottobock Quick Disconnect Wrist cannot be used with this wrist. For a prosthetic wrist to fit with a wide variety of devices, the wrist needs to be adaptable on both the proximal and distal ends. The proximal end needs to be adaptable to differing sockets and the distal end to various prosthetic hands. This modularity can be easily facilitated with 3D-printing.

#### Lightweight

Active, 2-degree-of-freedom prosthetic wrists that can be added to hands weigh between 95 to 500 grams [23]. As prosthetic device weight is correlated with abandonment, an active wrist should be as light as possible, certainly not exceeding 500 grams [4,27].

#### Functional

The device would need to be strong enough to lift external objects and perform tasks. One study showed that the average maximum wrist flexion and extension torques at neutral position for the human arm was 8.0 ± 3.0 N*m and 4.6 ± 1.0 N*m, respectively [28]. Ideally, a prosthetic wrist, would match this level of functionality. However, commercially available options are limited by space constraints, availability, and cost. As such, the motors must be as strong but small and lightweight.

### Device Design

A prosthetic wrist was designed in Solidworks (Solidworks, Waltham, MA) (Figure 1A) and subsequently assembled (Figure 1B). The body of the wrist consists of an interlocking rotation mechanism and cap. The interlocking rotation mechanism was designed to be 3D- printed with dissolvable support material, polyvinyl alcohol (PVA), between the interlocking parts. Once the support material is dissolved, the 2 subparts can freely rotate (Figure 1D). Each of these subparts houses a servo motor and the proximal subpart also houses an Adafruit Trinket M0 microcontroller (Adafruit Industries, New York, NY). The proximal servo motor is responsible for providing rotation between the 2 subparts. An Actobotics 525130 servo hub horn (RobotZone, Winfield, KS) is mounted to the proximal motor which is then fixed to the distal part via 4 screws. The distal servo motor is responsible for the additional degree of freedom (flexion/extension or radial/ulnar deviation) and has a servo hub horn mounted to it. The servo hub horn is fixed to the adaptable hand attachment portion of the wrist using 4 screws.

**Figure 1.**
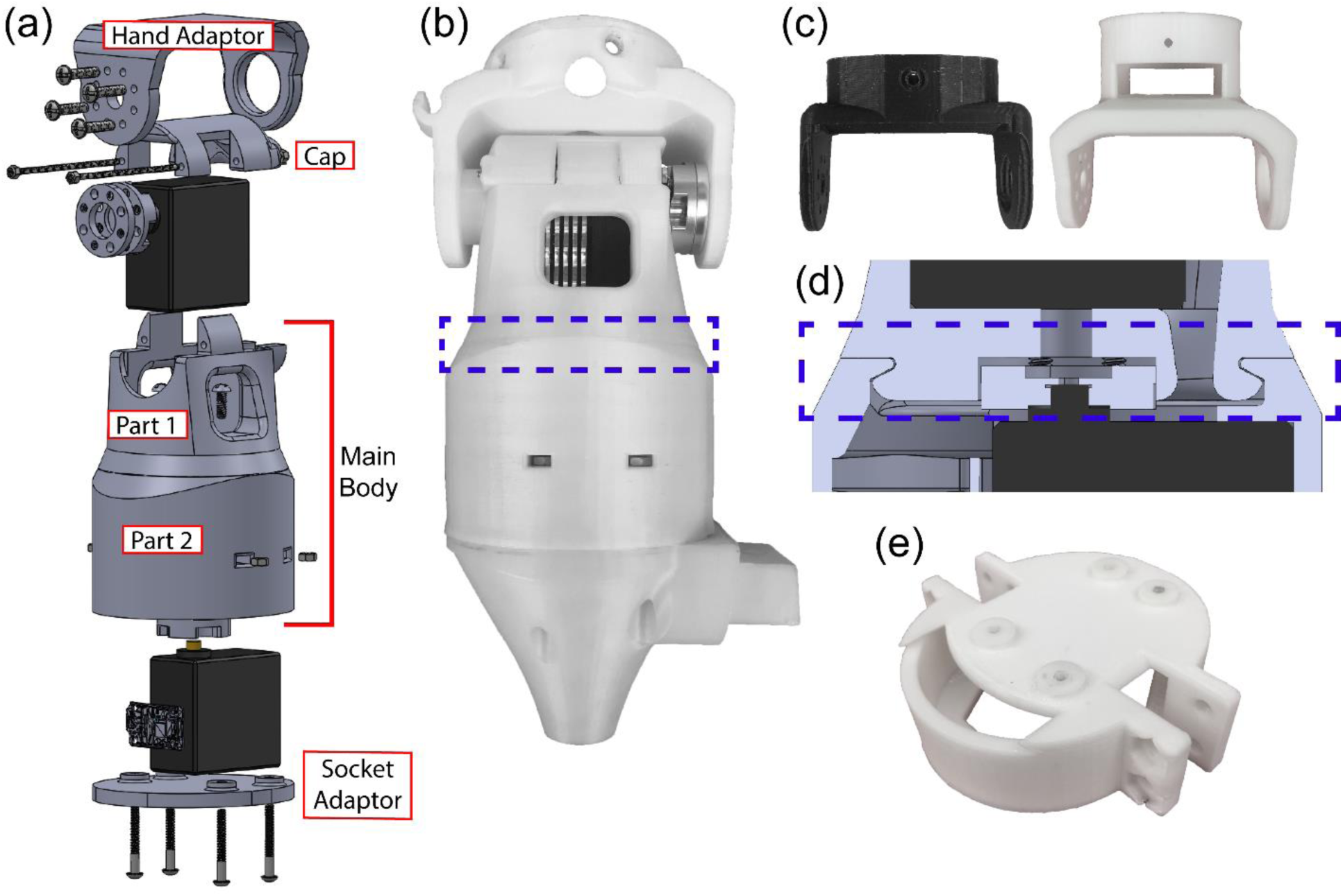
CAD design (a) and 3D-print (b) of the designed wrist. The hand adaptor (c) is easy to modify to fit to various prosthetic hands. The currently attached part (seen in b) is for the Ottobock quick disconnect. Attachments have also been made for the Ada [14] and Handi [16] hands. (d) The main body of the wrist contains subparts 1 and 2 which are 3D-printed with PLA and dissolvable PVA support material. These parts are interconnected via an interlocking rotational mechanism to preserve space. The proximal end can also be adapted to various sockets (e). Currently attached to the proximal end of the wrist in (b) is an attachment for the Bypass Socket [29].

Actobotics 545372 servo hub spacers (RobotZone, Winfield, KS) were used to provide proper spacing between the servo horn and the adaptable hand attachment portion of the wrist. The wrist can be manually adjusted to provide radial/ulnar deviation instead of flexion/extension. This is done by rotating the hand to the desired orientation before fixing it to the wrist. This modification changes the neural position of the hand 90 degrees. Because of this, reorienting the attachment of rotational servo motor the servo hub horn would be necessary.

#### Degrees of Freedom and Length

Because of the servo size and the length constraint of 12.5 cm, we could implement only a 2-degree-of-freedom wrist. Pronation/supination was included in the design because it is the most frequently implemented degree of freedom. The wrist was designed to be in series with either flexion/extension or radial/ulnar deviation: either can be selected by attaching the prosthetic hand of choice in the desired orientation. The length of the device without any modifications to the proximal and distal ends is 11.8 cm (Table 1).

**Table 1:**
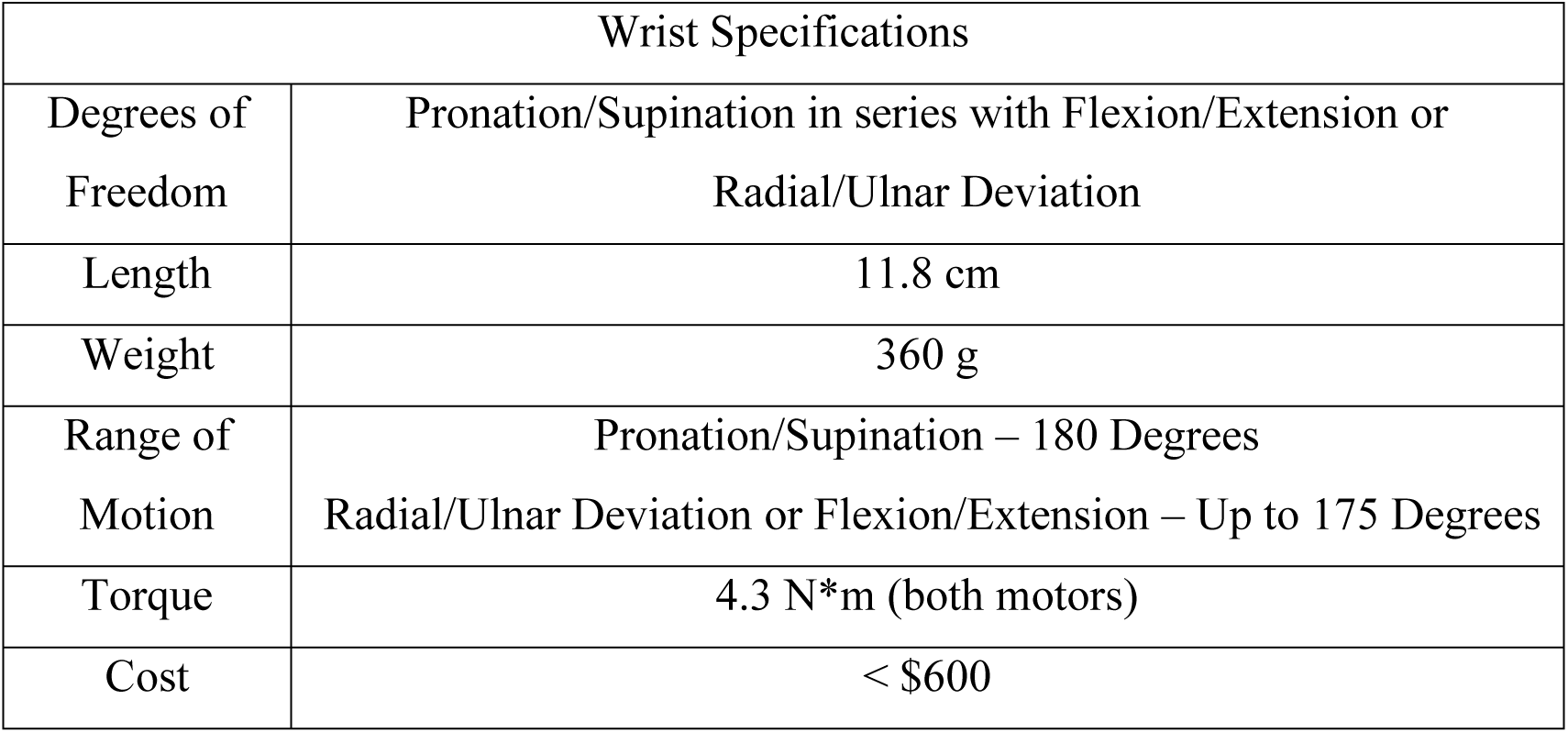
The wrist specifications fall within the established design criteria

#### Adaptable

The wrist accommodates virtually any hand and socket by using modular attachments. CAD design and 3D-printing can be used to allow for rapid adaptation to the distal and proximal ends of the device (Figure 1C and 3E). The distal hand adaptor was designed with a flat surface so modified attachments can be easily printed to fit new hands to the wrist. Likewise, the proximal end was designed to be easily modified to fit various sockets.

#### Lightweight

The device is 3D-printed with polylactic acid (PLA) which helps make the device inexpensive and lightweight—weighing only about 360 grams (Table 1).

#### Functional

High-powered hobby servo motors (Hitec D980TW, Hitec RCD USA, Poway, CA) were utilized due to their low cost and ease of use. This motor is a large, nonstandard sized hobby servo, and can provide up to 4.3 N*m torque, more than any other hobby servo readily available and closest to the healthy human wrist maximum extension torque of 4.6 ± 1.0 N*m [28] previously mentioned in the design criteria section. The two wrist motors are powered using a 7.5 V, 20 Amp power supply (967-CUS200LD7R5, TDK-Lambda Americas Inc., National City, CA).

### Assessment of Wrist Function

To understand the usefulness of an active wrist, we modified the clothespin relocation task: a simple task that relies on wrist function [30]. The experiment was completed by naïve, non-amputee participants (n = 8 participants) wearing a transradial bypass socket to simulate the experience of an amputee [29] (Figure 2). All participants gave written informed consent before taking part in experiments, in accordance with the University of Utah Institutional Review Board and the Department of Navy Human Research Protection Program. Participants were instructed to move a single clothespin from one location to another as fast as possible (See *Prosthetic Control Paradigm* section, below). If a participant dropped the clothespin, the trial was marked as a failure and discarded. Three variations of the task were completed (Figure 3A): a) moving the clothespin from a horizontal position to an adjacent horizontal position; b) moving the clothespin from a horizontal position to a vertical position (requiring wrist pronation/supination); c) reaching over a dowel to move the clothespin from a horizontal position to a horizontal position under the dowel (requiring wrist flexion/extension).

**Figure 2.**
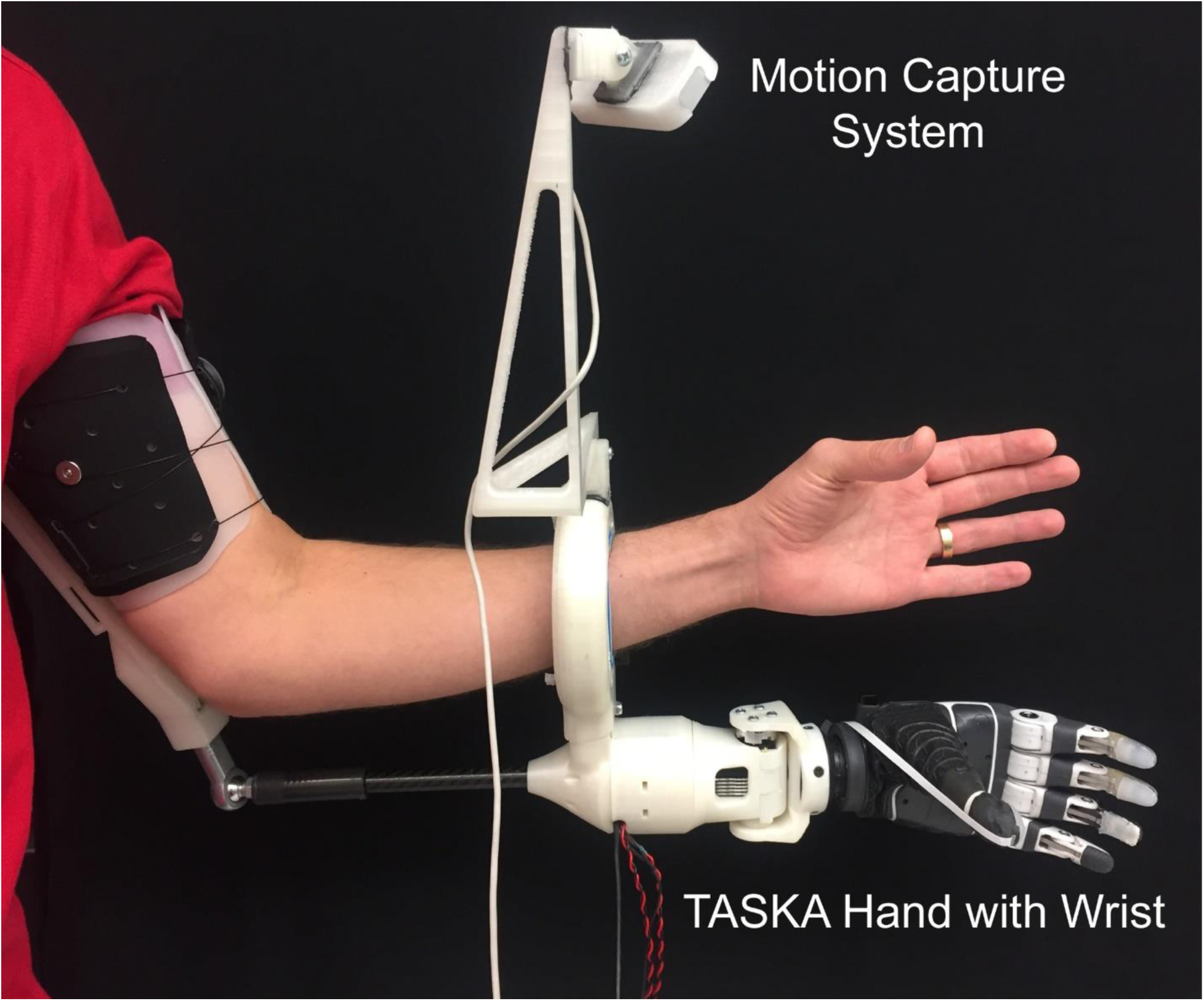
The intact participants wore a bypass socket [29] with a motion tracking system (Leap Motion) attached. Movement of the intact hand and wrist was tracked and translated into control of the prosthetic hand and wrist.

**Figure 3.**
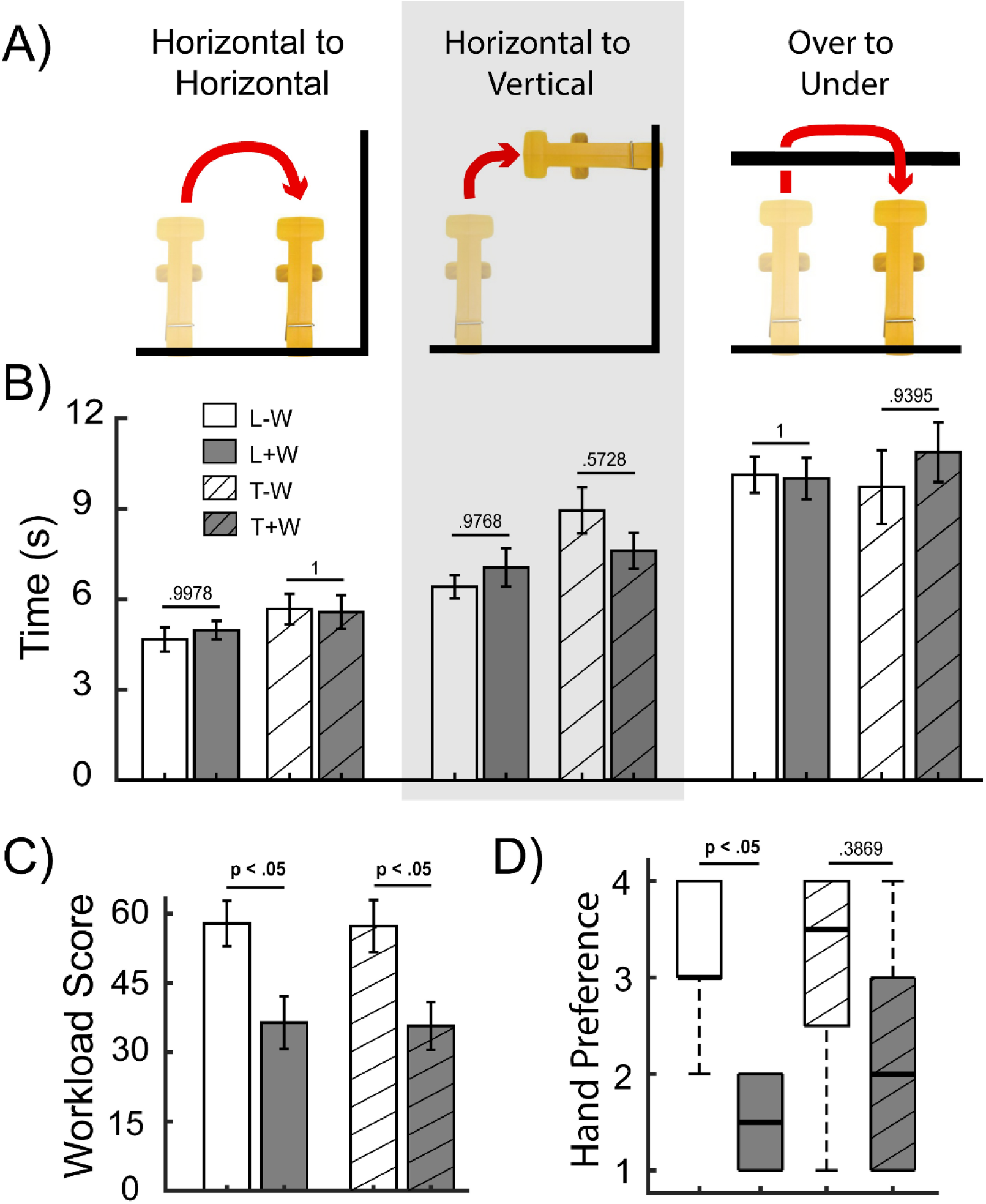
(A) The three variations of the clothes pin relocation task involved moving a clothespin from horizontal to horizontal position, horizontal to vertical position, and reaching over a horizontal bar to move the pin under the bar. (B) The implementation of a wrist did not significantly affect the time it took participants to complete tasks. (C) DEKA “LUKE” arm and TASKA workload scores were significantly reduced through the implementation of a wrist. (D) Arm preference as reported by participants for each condition. Bar plots show mean ± S.E.M. Box plots show median (center line), interquartile range (box), and most extreme non-outlier values (whiskers). *N* = 8 non-amputee participants. Significance differences were determined by a one-way ANOVA (or Kruskal-Wallis) followed by subsequent pairwise comparisons corrected for multiple comparisons. *p* values are only shown for primary comparisons (T+W vs. T-W, and L+W vs. L-W).

#### Prosthetic Control Paradigm

A motion capture system (Leap Motion, San Francisco, CA) was mounted to the top of the wrist rotation mechanism of the bypass socket (Figure 2) to track the position of the participants hands and wrist. Motion capture provided precise joint angles sampled at 60 Hz. Only a single degree of freedom (index finger flexion/extension) was provided to the hand to simplify the hand control and focus comparisons on wrist functionality. The index finger position of both hands was updated at 10 Hz. Due to inaccessible underlying software restrictions, the control of the DEKA “LUKE” Arm wrist differed from the wrist developed herein. The DEKA “LUKE” Arm required wrist position data to be converted to a velocity control signal. For the developed wrist, the joint angles of the wrist were averaged over a 1 second period to smooth the signal and used to control the wrist. For control conditions, both wrists were locked in a neutral position and was not actively updated with the motion tracking.

#### Experimental Conditions

Each participant completed the experiment under four conditions in a pseudorandomized counter-balanced crossover design, such that each participant served as their own control. The four experimental conditions were different hand/arm combinations, including: a) TASKA Hand with the 3D-printed wrist (T+W, first experimental condition); b) TASKA Hand with the 3D-printed wrist in locked position (T-W, first control); c) DEKA “LUKE” Arm with its embedded wrist fully functional (L+W, second experimental condition), and 4) DEKA “LUKE” Arm with its embedded wrist locked (L-W, second control). Our goal was to measure overall improvements associated with the wrist conditions (+W) relative to the no-wrist conditions (-W), such that the primary comparisons are T+W vs. T-W, and L+W vs. L-W.

The cross-over design was made up of 8 distinct experimental blocks performed one after another. Experimental blocks were pseudorandomized and counterbalanced to combat fatigue and motor learning effects. For each block, the participant performed the three tasks (horizontal-horizontal, horizontal-vertical, and over-under) four times each using a single hand-arm configuration. This resulted in a total of 8 trials per condition (2 blocks per condition, 4 trials per block). Participants were allotted up to 1 minute of practice before each block of trials. The median time of the 8 trials was reported as the participants final score.

#### Performance Metrics

We collected five performance metrics for each experimental condition: individual task completion time, workload across all tasks, and user preference across all tasks. The median time to completion (across the 8 trials) served as the completion time for each condition. After participants completed all tasks for a given experimental condition, workload was measured by having each participant complete the NASA Task Load Index survey [31]. The NASA Task workload score uses six workload-related factors to estimate an overall subjective workload score. At the end of the experiment, participants were asked to rank order the hand/wrist combination based on preference, such that a score of 1 is most preferable.

#### Statistical Analysis

All data were checked for normality using the Anderson-Darling and Lilliefors tests for normality. Outliers (more than 1.5 interquartile ranges above the upper quartile or below the lower quartile) were removed prior to statistical analyses. A one-way analysis of variance (ANOVA), or non-parametric equivalent (Kruskal-Wallis), was then performed as appropriate across the four experimental conditions per performance metric. If any statistical difference was found, subsequent pair-wise comparisons were performed using the Dunn-Sidak correction for multiple comparisons.

## RESULTS

### Wrist Performance and Assessmente

There were no statistically reliable differences in time to completion for hands with functional wrists compared without functional wrists. However, providing a wrist reduced subjective workload for both the LUKE and TASKA hands, as measured on the NASA Task Load Index. Subjects also preferred the LUKE hand with a wrist, over a LUKE hand without a wrist.

### Horizontal to Horizontal Task

The average time for the participants to complete the task was 4.67 ± 0.40 for the L-W, 4.97 ± 0.31 for the L+W case, 5.67 ± 0.51 for the T-W, and 5.57 ± 0.56 for the T+W case (Figure 3B). The one-way ANOVA showed no significant differences among the four experimental conditions (*p* = 0.355).

#### Horizontal to Vertical Task

The average time for the participants to complete the task was 6.41 ± 0.39 for the L-W case, 7.05 ± 0.63 for the L+W case, 8.94 ± 0.76 for the T-W case, and 7.60 ± 0.60 for the T+W case (Figure 3B). The one-way ANOVA showed significant differences among the four experimental conditions (*p* < 0.05). We found no significant differences among the primary comparisons of interest (wrist + vs. wrist -); however, significant difference was found between T-W vs. L-W (*p* = 0.039).

#### Over to Under Task

The average time for the participants to complete the task was 10.12 ± 0.59 for the L-W, 10.00 ± 0.69 for the L+W case, 9.71 ± 1.22 for the T-W case, and 10.88 ± 0.99 for the T+W case (Figure 3B). The one-way ANOVA showed no significant differences among the four experimental conditions (*p* = 0.827).

#### NASA Task Load Index Scores

Functional wrists decreased subjects’ subjective workload as measured by the NASA Task Load Index. The overall workload required to use the various hand/wrist combinations was 57.88 ± 4.94 for the L-W case, 36.42 ± 5.69 for the L+W case, 57.33 ± 5.67 for the T-W case, and 35.71 ± 5.16 for the T+W case (Figure 3C). The one-way ANOVA showed significant differences among the four experimental conditions (p < 0.05). Significant differences were found in both primary comparisons of interest (L+W vs. L-W and T+W vs. T-W). The workload was significantly less for the L+W vs. L-W (*p* = 0.0026) as well as for the T+W vs. T-W (*p* = 0.0024). Secondary comparisons also showed a significant difference in T+W vs. L-W (*p* = 0.0018) and T-W vs. L+W (*p* = 0.0033).

#### Subjective Hand Preference

The L+W was selected first 4 times, second 4 times, third 0 times, and fourth 0 times. The T+W was selected first 3 times, second 2 times, third 2 times, and fourth 1 time. The T-W was selected first 1 time, second 1 time, third 2 times, and fourth 4 times. The L-W was selected first 0 times, second 1 time, third 4 times, and fourth 3 times. The average choice of each hand combination was: 1.5 for L+W, 2.125 for T+W, 3.125 for T-W, and 3.25 for L-W (Figure 3D). The Kruskal-Wallis showed statistical difference between the L+W vs. L-W (*p* = .0123). No statistical difference was found between the T+W vs. T-W (*p = .3869).* The lack of statistical difference in the T+W vs. T-W case could be due to the n=8 sample size. A secondary comparison between the L+W vs T-W also showed statistical difference (*p* = .0251).

#### Compensation

Figure 4 shows two individuals performing the horizontal to vertical and over to under tasks. We observed that during the conditions without a wrist, the participants compensated by rotating their shoulders and torso.

**Figure 4.**
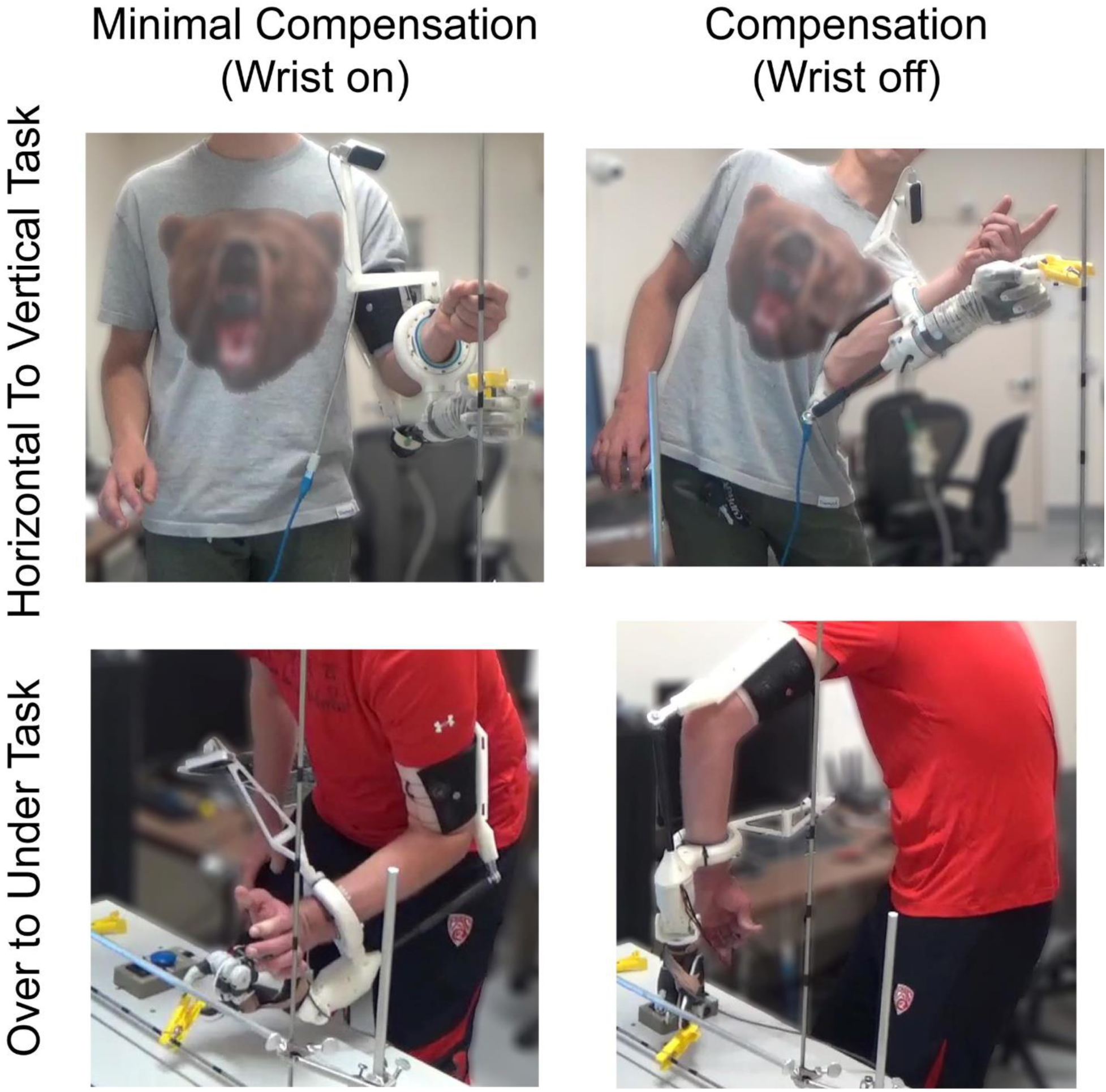
Participants relied on compensation to complete tasks that otherwise would require a wrist.

## DISCUSSION

We developed a 3D-printable, lightweight, and adaptable prosthetic wrist, which met or exceeded our design criteria (Table 1) and demonstrated that our wrist (as well as the DEKA “LUKE” Arm’s wrist) significantly reduced workload as measured by the NASA Task Load Index in a modified clothespin relocation task. Participants preferred a wrist over no wrist for the L+W vs L-W comparison (*p* = .0123). A larger sample size may be necessary to reveal significance between the T+W vs T-W condition. Although not a primary comparison of interest, for all 5 of the performance metrics, there were no significant differences between the L+W and T+W. This suggests that our wrist is not significantly worse than the commercially available DEKA “LUKE” Arm for these 5 performance metrics..

### Implementation of a Wrist Significantly Reduces Workload but Does Not Affect Speed

Although having a wrist did not affect the time it takes for a participant to complete these particular tasks (Figure 3b), it did reduce the workload required for these tasks (Figure 3c). Although we did not explicitly quantify compensatory movements, we visually observed that participants heavily used compensatory motions to complete the tasks when a wrist was not available (Figure 4). Anecdotally, and without explicit measurement, the addition of the wrist reduced compensatory movements (Figure 4). It is likely that the NASA Task Load index scores for workload were higher for the no-wrist cases due to the physical exertion required to use compensatory motions that the participant needed to perform to complete the tasks.

For timed tasks, statistical difference was only found in the time it took to complete the horizontal to vertical task with the T-W vs. L-W, which suggests that performance of the two hands differ. This could be due to differences in weight, grip strength, grip material or hand properties. For example, the DEKA “LUKE” Arm flexion and extension are coupled with radial and ulnar deviation. With the addition of the wrist, differences between the DEKA “LUKE” Arm vs. the TASKA hand were no longer present, suggesting that the wrist may preferentially benefit less functional hands. Comparing the T+W and L+W conditions was not our primary goal, but we observed no statistical differences among these conditions.

After the experiment, participants were asked to rank their preference of the prosthetic in order from 1-4 (Figure 3d). Collapsing across the two prosthetics, the experimental conditions without a functional wrist required a greater workload (*p* < 0.05, paired t-test) and were preferred less (*p* < 0.05, Wilcoxon rank-sum test). Together, these results suggest that, despite there being no significant improvement in the time it took the participants to complete tasks, the addition of a functional wrist is preferable because it reduces the subjective workload needed to complete tasks.

### Other Developed Wrists Have Limitations in Length or Strength

Other 1-, 2-, and 3-degree-of-freedom active prosthetic wrists have been developed. However, few of these wrists can be 3D-printed and/or provide as much torque [22,23] (Table 2). One 2-degree-of-freedom wrist uses a differential system to conserve space. At just 4.8 cm, the wrist is very short; however, it provides just 0.073 N*m of torque for wrist flexion [32], which would not lift most prosthetic hands. The wrist discussed in this paper, though longer, uses motors that are much more practical and that can provide roughly 60 times more torque (4.3 N*m), which is much more realistic for of daily living.

**Table 2:** Adapted from Bajaj Et. Al [23]to include only active wrists with more than 1 degree-of-freedom with developed wrist

| Product Name/Lead<br>Author | DOF | 3D Printed | Commercial | Diameter<br>(cm) | Length<br>(cm) | Weight<br>(g) | Stall Torque<br>(N*mm) |
| --- | --- | --- | --- | --- | --- | --- | --- |
| *Olsen, N. R.* | 2 | Y | N | 7.4 | 11.8 | 330 | 4300 |
| Abd Razak, N. A. [34] | 2 | N | N | -- | -- | 500 | 13000 |
| Controzzi, M. [35] | 2 | N | N | -- | -- | 240 | -- |
| Kyberd, P. J. [32] | 2 | N | N | 9.6 | 5.0 | 200 | 73 |
| Roose, C. [36] | 2 | Y | N | 5.3 | -- | 95 | 321(flexion) |
| RIC Arm [37] | 2 | N | N | -- | -- | 170/155 | 900/1000 |
| RSL Stepper bebionic<br>Wrist [38] | 2 | N | Y | 5.0 | 7.5 | -- | -- |
| Mahmoud, R. [39] | 3 | Y | N | - | - | - | 930/210/210 |
| Dange, S.V. (24) | 3 | Y | N | 10.16 | 26.67 | -- | 4300 |

Another wrist, developed by Dange [33], was capable of three degrees of freedom. This wrist is more than double the length of our wrist, making the wrist the length of the radius bone itself. The Dange wrist uses servo motors that provide the same torque and are the same dimensions as the ones used by our wrist. However, our wrist is far more compact and realistic as a prosthetic device, although limited to only 2 degrees of freedom.

### Limitations and Future Work

Due to the relatively limited availability of amputees, the experiment was performed on only intact participants using a bypass socket [29]. Paskett et al. showed that the bypass socket can closely simulate the experience of an amputee; however, future studies should broaden the testing of the wrist to include amputees.

One limitation of the servo motors is that extensive use at maximum load produces heat capable of softening the PLA. Future work would perform tests to quantify the extent of this overheating and could include efforts to reduce overheating by replacing servos with brushless motors (which draw less current and therefore create less heat), using a more heat resistant 3D- printing material, or adding an explicit heat sink and cooling vents.

Our wrist can be used in many research applications. For example, one study showed that a single-degree-of-freedom prosthetic hand with a wrist may be comparable to a dexterous hand without a wrist (10). However, this study involved the use of an intact person with a brace on their arm limiting certain movements. Our wrist could extend this study and allow experimentation with a variety of prosthetic devices instead of an intact hand. The adaptable aspect of our wrist also enables it to be used to study optimal control algorithms for prosthetic wrists.

## CONCLUSION

Our novel 3D printed wrist can be used with many terminal devices and sockets to reduce task load. The wrist can also be used in research to show how prosthetic devices can be improved with the implementation of a functional wrist. Ultimately the wrist may aid in the development of more biomimetic prosthetic devices and improve the quality of life for amputees.

## Acknowledgments

We thank TASKA Prosthetics for the use of the TASKA hand in this study. We likewise thank DEKA for the use of the “LUKE” arm. We would also like to acknowledge the University of Utah Center for Medical Innovation for assisting us with the 3D printing of the wrist.

## Funding

This work was sponsored by the Hand Proprioception and Touch Interfaces (HAPTIX) program administered by the Biological Technologies Office (BTO) of the Defense Advanced Research Projects Agency (DARPA), through the Space and Naval Warfare Systems Center (contract no. N66001-15-C-4017). Additional sponsorship was provided by the NSF through grant no. NSF ECCS-1533649 and NSF GRFP award no. 1747505.

## Author contributions

N.R.O. designed and built adaptable wrist, developed software for the wrist, designed the experiment, ran the experiments, 3D printed components for the experiment, performed statistical analysis, and drafted the manuscript.

J.A.G. provided design oversight, developed software to control the prosthetics, designed the experiment, provided guidance on statistical methods, and drafted the manuscript. M.R.B. designed the experiment, provided guidance on statistical methods, and drafted the manuscript.

M.D.P provided wrist design oversight, developed the bypass socket used in the experiment, and provided guidance on statistical methods.

D.T.K. provided wrist design oversight, and designed software to control the prosthetics.

T.N.T. developed software to control the prosthetics.

C.C.D. provided clinical support, oversights, and relevance throughout.

G.A.C. oversaw and led the development of all methods, experiments, and protocols and assisted with experiments and drafting of the manuscript.

All authors contributed to the revision of the manuscript. The views expressed herein are the authors’ and do not necessarily reflect the views of their employers or funding sources.

## Competing interests

D.T.K. is now an employee of Blackrock Microsystems.

## Data and materials availability

Deidentified data will be made available upon request. Data and materials requests should be sent to G.A.C. Distribution of identifiable data for human subjects will require approval of the University of Utah Institutional Review Board.

